# Real-time Biomechanical Characterisation of Cytoskeletal Remodelling

**DOI:** 10.1101/2024.05.29.595860

**Authors:** Kaiwen Zhang, Chayla Reeves, Joseph D. Berry, Kate Fox, Aaron Elbourne, Amy Gelmi

## Abstract

As progenitors for tissue, human mesenchymal stem cells (hMSCs) with ability of self-proliferation and differentiation into various cell types such as osteocytes and adipocytes show great potential applications for tissue engineering. Stem cell fate regulation is highly affected by the cytoskeleton structure and mechanical properties. In this paper, quantitative Atomic Force Microscopy (Q-AFM) was used to continuously characterise topography and biomechanical properties while applying cytoskeleton disruptors to hMSCs. The cell stiffness (quantified by Young’s modulus), primarily governed by the cytoskeleton network, had quantifiable changes associated with cytoskeleton polymerisation and depolymerisation when treatments were applied. Furthermore, with Q-AFM measurements, these changes were tracked in real time over a period of minutes to hours, and the biomechanical properties of the cells were tracked through the applied treatment and subsequent recovery post treatment. Here we present the capability of Q-AFM to perform real time biomechanical characterisation of living cells, directly correlated to intracellular structure and cytoskeletal remodelling.

## Introduction

Human mesenchymal stem/stromal cells (hMSCs) have become a popular cell source for stem cell-based tissue engineering therapy. The self-renewing ability and multipotent differentiation ability into cell lineages such as osteocytes, adipocytes and even neurons bring up the uniqueness of hMSCs.^1–6^ hMSCs derived from patients are a promising avenue for autologous tissue engineering, which offers advantages such as reduced need for immunosuppressive drugs, and lower risk of host vs graft disease during transplantation. Potential applications of hMSCs in regenerative medicine have been reported, such as bone and cartilage repair, and skin regeneration.^7–12^

The cytoskeleton, a network of filaments and tubules that extends throughout the cell, not only supports cell morphology, adhesion, and migration, but also plays an important role in cell signalling and potential cell fate decision.^13,14^ The cytoskeleton is the main component contributing to cell stiffness, with its ability to resist deformation caused by external force while it can also be dynamically regulated to adapt to the extracellular matrix.^15^ Seeding cells on substrates mimicking tissue-like stiffness, cells sense and respond to the substrate stiffness including cytoskeleton realignment and cell stiffness adjustment, promoting to differentiation into a specialised cell phenotype. For example, substrates with stiffnesses ranging from 0.1-1 kPa (brain tissue), 8-17 kPa (muscle tissue), 25-40 kPa (bone collage) induce neurogenesis, myogenesis, and osteogenesis, respectively, with diffuse F-actin on soft gels to progressively organised F-actin on stiffer substrates.^16^ Under growth factor induced differentiation, hMSCs became 1.6-fold stiffer during osteogenesis along with thicker cytoskeleton network,^17^ while the Young’s modulus of hMSCs decreased from 3.2 ± 2.2 to 0.9 ± 0.8 kPa during adipogenesis.^18^ It is possible to predict cell fate decision by monitoring cellular mechanics with external stimulus. Disruption of the cytoskeleton has also been reported to affect cell fate via specific cell signalling pathways. However, the dynamic response of cells to the disruptors hasn’t been understood from previous studies, especially changes to the cytoskeleton organisation and cellular mechanics on the same cell.

Mechanotransduction stands as one of the fundamental mechanisms through which stem cells perceive and react to external mechanical stimuli. When cells sense physical stimuli via membrane components like integrins, mechanotransduction pathways are activated, leading to the conversion of mechanical signals into biochemical cues that can be interpreted by intracellular features. These signals are then transmitted to the nucleus, influencing gene expression via the cytoskeletal network. Simultaneously, the cytoskeleton undergoes reorganisation, depolymerisation, and polymerisation in response to the signals, significantly impacting cell stiffness, which in turn affects cellular fate.^19,20^ Recent investigations into mechanotransduction processes have primarily concentrated on biomolecular analyses, often employing fluorescent imaging and PCR techniques.^21^ Traction force microscopy (TFM) can be used to measure intracellular tension,^22^ but this technique is limited to transparent substrates embedded with fluorescent tracer beads. Combined with AFM, TFM has been used to correlate contractile forces with biomechanical properties of cells.^23^ However, there has been a significant gap in exploring how alterations in cytoskeletal structure influence both cell stiffness and cellular responses.^24,25^ It is crucial to monitor changes in the mechanical properties of the cytoskeletal network during mechanotransduction processes. Such analysis is essential for unravelling the complex interplay between mechanical stimuli and cellular responses, ultimately regulating various cellular processes, including cell adhesion, migration, differentiation, and tissue morphogenesis.

The quantitative Atomic Force Microscopy (Q-AFM) was developed in 2013,^26^ making it possible to measure cell mechanics non-invasively and gently with high resolution in a short time frame, achieving temporal analysis of single cells with external treatment. The scanning frequency of Q-AFM measurements can reach at least 1 kHz, while the sampling rate of the traditional force-volume mapping mode is around 1-200 Hz, which means the advanced Q-AFM measurement is much faster and provides higher resolution.^17,18,27,28^ The large set of force spectroscopy measurements obtained at each pixel allows correlative topographical images, stiffness images, and adhesion images to be generated simultaneously.

In this paper, we demonstrated Q-AFM to monitor cellular mechanics on single cells in real-time with cytoskeleton disruptor treatments. The dynamic change of the cytoskeletal structure and cell mechanics were correlated with the AFM measurements. Confocal microscopy was used to verify the observation of the cytoskeleton structure. With the ability to monitor cell stiffness in real-time, the mechanotransduction process can be studied in the future while applying various external stimulations to stem cells to affect their cell fate.

## Method

### Cell culture

Bone marrow derived hMSC (Lonza, Switzerland) were cultured and maintained at 37 °C and humidified atmosphere with 5% CO_2_ in alpha Minimum Essential Media (αMEM, Gibco, USA), 1% Penicillin-Streptomycin (Gibco, USA), and 10% FBS (Gibco, USA) until 80% confluency. For the chemical treatment experiments, cells were seeded on 35 mm FluoroDishes (WPI, USA) at 1000 cells/cm^2^ seeding density. hMSCs up to passage 6 were used.

### Cytoskeleton Disruption

Cytochalasin D (CytD, 5 mg/mL in DMSO, Sigma, USA) was added to CO_2_ independent Leibovitz’s L15 media (Gibco, USA) to reach concentrations of 0.1, 0.5, and 1 µg/mL. Blebbistain (Sigma, USA) was dissolved in DMSO to reach a concentration of 1.6 mM as stock solution. Stock solution was diluted with L15 media to obtain concentrations of 12.5, 25, 50 µM for hMSCs treatment. Colchicine (Sigma, USA) was dissolved in Ethanol with 0.876 mM as stock solution, which was diluted with L15 media to concentrations of 0.5, 5, 50 µM for cell treatment. Nocodazole (Sigma, USA) was dissolved in DMSO as stock solution with a concentration of 2 mg/mL, which was also diluted to 1, 5, 10 µg/mL with L15 media.

During AFM measurements, cells were exposed with treatment solutions ranging from 30 to 120 min, depending on the response speed of cells towards the cytoskeleton disruptor. Once the cells showed obvious structural changes, treatment solutions were replaced with fresh L15 media and measurements continued for 120 min. At least 5 cells for each chemical disruption were characterised through the whole period of treatment and removal, 2 with higher resolution showing clear intracellular structure and 3 with relatively lower resolution for monitoring dynamic of whole cell stiffness.

### Cytoskeleton staining

Cells were fixed in formaldehyde at five time points; before treatment, 1h of treatment, 2h of treatment, 1h post treatment, and 2h post treatment. ActinRed™ 555 ReadyProbes™ Reagent (Invitrogen, USA) or ActinGreen™ 488 ReadyProbes™ Reagent (Invitrogen, USA) and NucBlue™ Live ReadyProbes™ Reagent (Invitrogen, USA) were added to the cell dishes with 2 drops/mL to stain the cytoskeleton and nuclei respectively for 20 min. Cells were washed with PBS (Gibco, USA) and imaged by widefield microscopy (Olympus, Japan) and confocal microscopy (Nikon Ti-E, Japan). The images were analysed using Fiji ImageJ.

### Atomic force microscopy

Nanowizard IV (JPK instruments, Germany) Q-AFM was used for cell structure and mechanics monitoring. Measurements were conducted in L15 media with the temperature controlled at 37°C in a PetriDishHeater (Bruker, Germany), using ContGD-G cantilever (BudgetSensors, Bulgaria) with spring constant 0.2 N/m.

Customised programming in Python was used to fit all the force-distance curves with the Hertz model^29^ using a parabolic indentor, with fitting range less than 18% of cell thickness,^30^ to obtain Young’s modulus of hMSCs. Contact points were estimated using the zero-gradient method.^31^ The relationship between force F, cell indentation δ, probe tip radius R, Poisson’s ratio υ and Young’s modulus E of Hertz model, is described as below. The substrate corrected model developed by Garcia, P. D., & Garcia, R.^32^ and the comparison between substrate uncorrected (basic Hertz model) and corrected model are shown in S1.

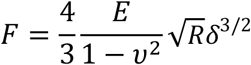

### Statistics

Significance was determined at p ≤ 0.05. Fitted Young’s modulus were analysed with a one-way analysis of variance between before, during and after exposure to cytoskeleton disruptor. All statistical analysis was carried out using GraphPad Prism.

## Result and discussion

### Correlating AFM measurement and confocal microscopy imaging

While AFM is primarily used for surface characterisation of materials, measuring the intracellular structure of biological samples, especially for cells, is achievable with the Q-AFM, with its working mechanisms shown in Fig 1a. The live hMSCs were imaged by Q-AFM, which provides highly resolved topographical and biomechanical information through analysis of each individual force-distance curve. The force-distance curves were fitted to determine the Young’s modulus, illustrating correlative topographical features with their corresponding stiffness. To confirm features observed in the AFM images represent the cytoskeletal structure of the hMSC, namely the actin structures, live hMSC were scanned by Q-AFM and subsequently fixed and prepared for immunofluorescent imaging.

**Figure 1.**
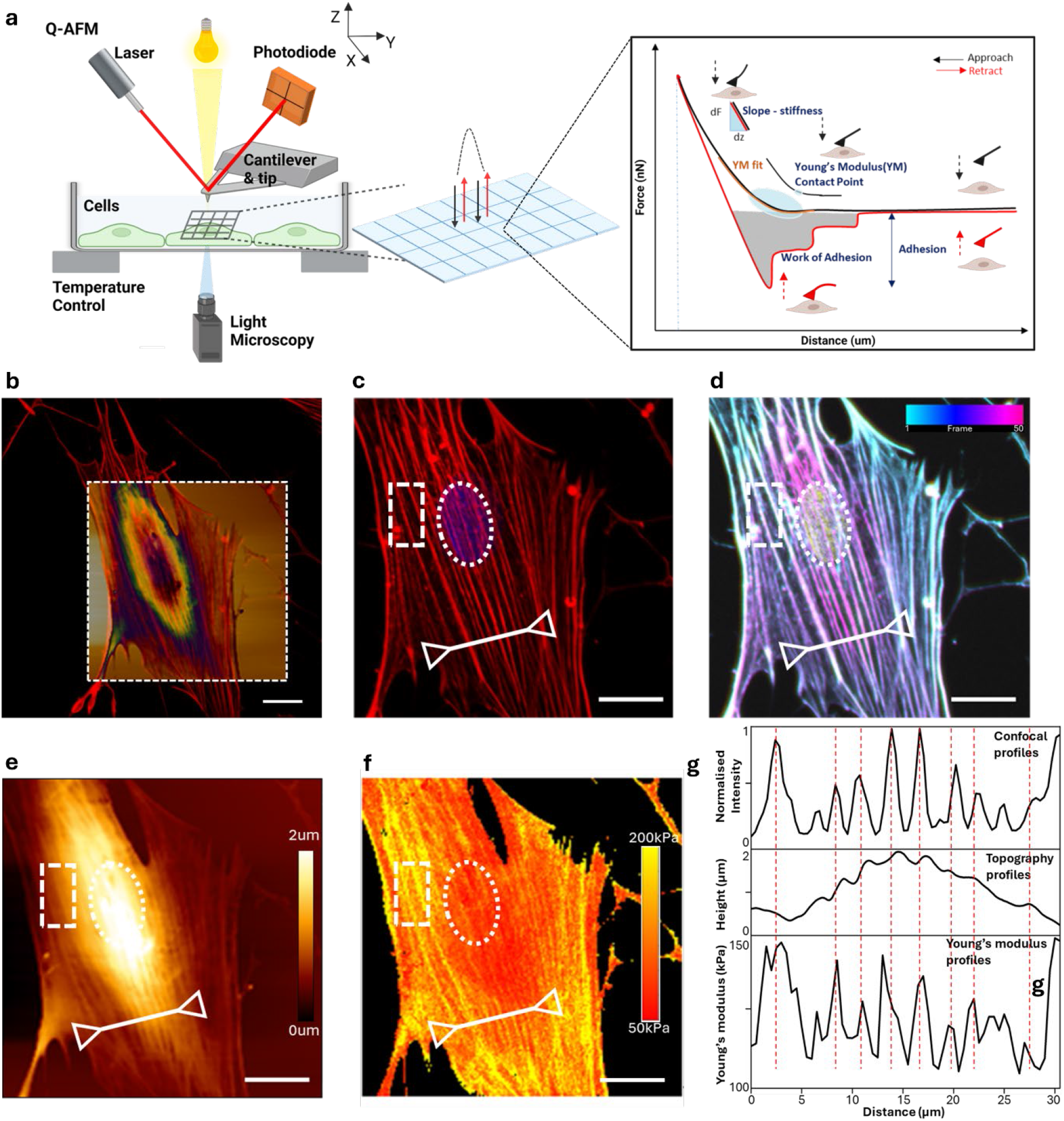
Correlating Q-AFM measurements with confocal images. (a) Schematic of Q-AFM working mechanisms, (b) Q-AFM topographical image stacking on confocal image to obtain the same region for comparison, (c) confocal image (actin stains), (d) Z-stack confocal image with colour scale frame 1-50 at the top right corner, (e) Q-AFM topographical image, (f) Q-AFM Young’s modulus map obtained by fitting force-distance curves, (g) profiles of the line with arrows drawn in pictures c-f. Scale bar is 20 µm.

In Fig 1c, the actin and nucleus are clearly visualised, and a depth-coded image reveals the apical and basal actin features (Fig. 1d). The corresponding Q-AFM scan of the hMSC reveals topographical features correlating to the apical actin (Fig. 1e), with the overall cell size shrinking around 5% post fixation. Cell morphology assessed using both techniques exhibited remarkable consistency, presented in the stacked images in Fig 1b. The cytoskeleton structure characterised by confocal images appeared relatively distinct, which in turn are present in the Q-AFM image.

The corresponding Young’s modulus map is shown in Fig 1f, determined via fitting of more than 30,000 individual force-distance curves. There are clear areas where the Young’s modulus is higher (stronger yellow signal) which correlate with the actin features observed in the topographical scan and confocal images. The force-distance curves have been fitted to an indentation depth 200 - 300 nm, while the cell membrane thickness is nominally less than 10 nm.^33,34^ Hence, it is reasonable to assume that we are measuring the stiffness of the apical actin structures within the living cell. In between the actin structures, we can assume that we are measuring the cell membrane and cytosol.

Further correlating the intracellular features across confocal images, Q-AFM topographical images, and Young’s modulus maps, profiles of the same line (marked by a white line with arrows) from these images were extracted and plotted in Fig 1g. The peaks in fluorescent intensity corresponded to the presence of individual actin fibres, which also correlated with peaks in the Young’s modulus profiles. This indicated that the actin was significantly stiffer than other intracellular structures, and the stiff structure observed in Young’s modulus were parts of the actin cytoskeleton. In the topographical profiles, which measured the height of the cell body, the results pertaining to actin did not return to baseline, showing no peaks for the induvial actin but small steps along the curves, which largely aligned with peaks in the other two profiles and the cytoskeletal features in the topographical images.

Referring to the Young’s modulus map, the stiffer part (yellow in colour) was not always distributed at the cell edge, indicating minimal substrate effect during data fitting. Cell stiffness was heterogenous within the cell, depending on the distribution of fibre features and the nucleus. When comparing the stiffness between the nuclear and perinuclear regions, represented as white dots in circles and squares respectively in Fig 1e, the perinuclear region, featuring a thick actin bundle, exhibited 1.3-fold stiffer than the nuclear region. This suggests that the cytoskeleton is stiffer than the nucleus, thus predominantly contributing to cell stiffness. However, the measurement of nuclear region is also influenced by the nucleus’ location within the cell. If the nucleus is positioned apically, the Q-AFM mainly characterises the stiffness of nucleus, whereas when the nucleus is close to the substrate, the structure above nucleus is measured, and its stiffness is dependent on the distribution of actin. It is notable that the cell stiffness measured by Q-AFM in this study is considerably higher than that reported in previous literature (less than 5 kPa)^18,35,36^, mainly attributed to the sharpness of the tip used. Research utilising Q-AFM with tips as small as 10 nm in radius has also shown significantly elevated cell stiffness, with Young’s modulus reaching up to 100 kPa for *NIH3T3* fibroblasts and 600 kPa for *PaR3* cells.^37,38^ It is worth mentioning that using sharp tip is more sensitive to local variations in cell structure and hence stiffness than a large spherical probe, which averages out stiffness over a larger area.^39^

The characteristics revealed in Q-AFM topographical images exhibited a strong correlation with those observed in confocal images of identical cells, aligning with the structural features illustrated in the Young’s modulus maps. This indicates that the primary features captured in Q-AFM measurements represent the cytoskeleton, which is the principal contributor to cell stiffness. Consequently, Q-AFM holds promise for real-time monitoring of intracellular structures, particularly the cytoskeleton, and quantification of cell mechanics of the same cells.

### Biomechanically characterising the effect of cytoskeletal disruptors on cell structure

Q-AFM is capable of characterising dynamic cytoskeleton remodelling of a cell. The cytoskeleton disruption chemicals are known to affect different cytoskeleton components, with their binding target and effect shown in Table 1. To evaluate Q-AFM as a tool to track and quantify cytoskeletal remodelling in real-time, hMSCs were treated with cytoskeleton disruptors to induce cytoskeleton remodelling.

**Table 1.** Effect of cytoskeleton disruptors on intracellular features of hMSCs.

| Chemical | Target | Effect | Selected concentration |
| --- | --- | --- | --- |
| CytD | Actin filament | Inhibiting actin polymerisation | 1 µg/mL |
| Blebbistatin | Myosin II | Inhibiting contractility of actomyosin via blocking myosin II ATPase activity | 12.5 µM |
| Nocodazole | Microtubule | Inhibiting microtubule polymerisation | 5 µg/mL |
| Colchicine | Tubulin | Inhibiting microtubule polymerisation, promoting actin polymerisation | 5 µM |

Based on previous literature,^17,40–43^ preliminary experiments to determine the suitable concentrations for cell treatment were conducted, where changes in cytoskeletal structure could be observed while cell viability is maintained, and a cellular response happens within a reasonable time frame (less than 2h). The actin cytoskeleton of the treated hMSCs were assessed using fluorescent imaging (Fig S2), and the selected concentration for each disrupter is listed in Table 1.

The representative results of all four treatments are shown in Fig 2. The disruption to the cytoskeleton from the chemicals is evident through the actin staining of fixed hMSCs, post-treatment (Fig 2 subset ii), comparing to the pre-treatment with clear actin figures shown in Fig 2 subset i. The AFM topographical images illustrate the change to the cytoskeletal structure in real-time as the AFM scanning was performed on the same live cells, where a scan of the hMSC pre- and post-treatment were generated (Fig 2 subset iii and iv respectively). From the Q-AFM data, the Young’s modulus for each point is also quantified (Fig 2 subset v and vi).

**Figure 2.**
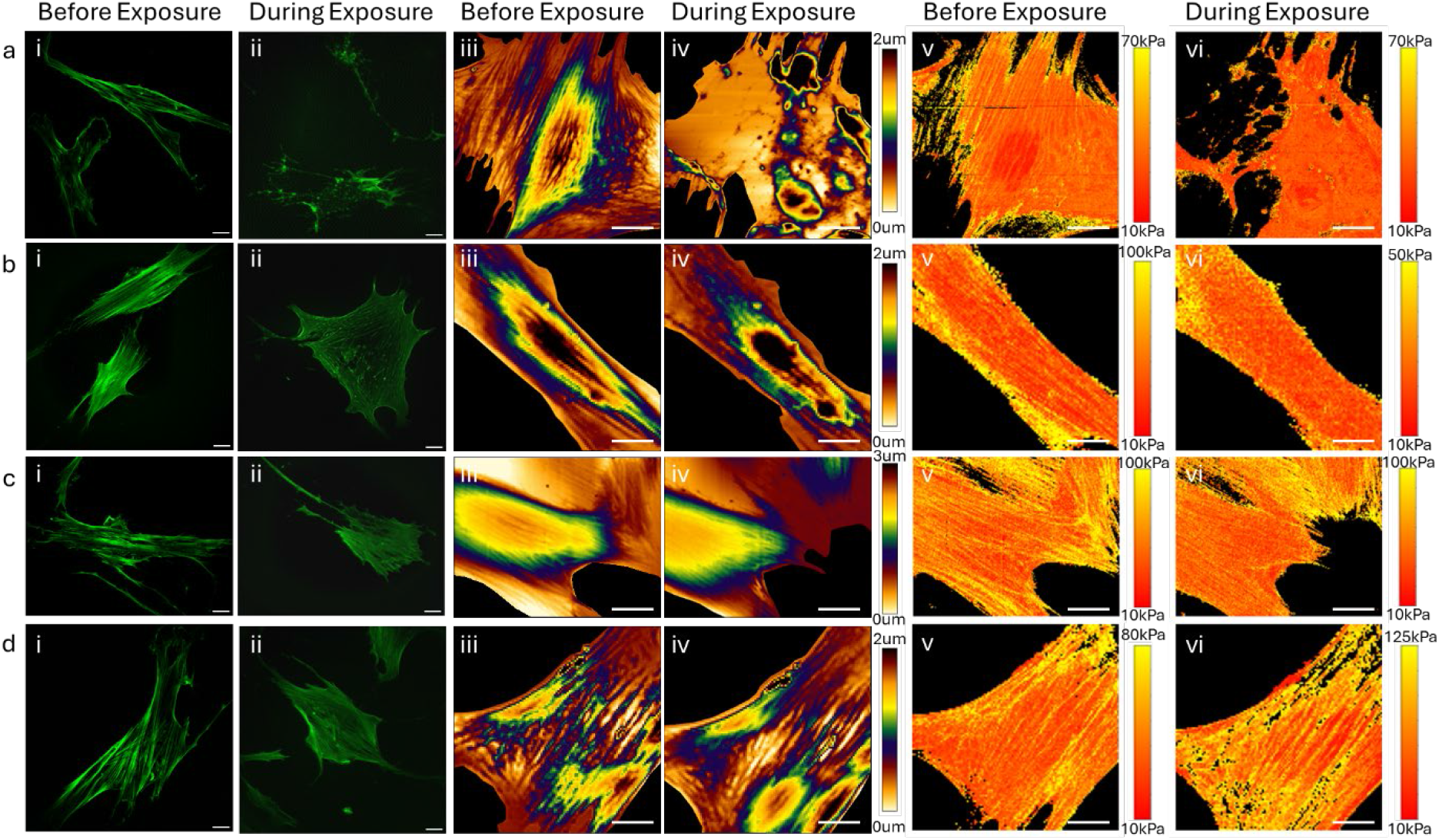
Representative results of cytoskeleton disruptors treating hMSCs, namely 1 µg/mL CytD (row a), 12.5 µM blebbistain (row b), 5 µg/mL nocodazole (row c) and 5 µM colchicine (row d). Subset i and ii are fluorescent images (actin stains) of before- and during exposure captured by widefield microscopy. Subset iii-iv are topographical images measured by Q-AFM. Subset v-vi are Young’s modulus maps obtained by fitting the F-D curves of Q-AFM. Scale bar is 20 µm Timeframe of each AFM measurement was 20 min.

Cells treated with CytD, blebbistatin or nocodazole treatment showed depolymerised cytoskeletal structure, shown in Fig 2 row a-c, illustrating a decrease in Young’s modulus. Specifically, the effect of CytD was more significant, with most of the cytoskeleton structure being degraded, observed in both fluorescent images and AFM topographical images, with 41% decrease in Young’s modulus. In comparison, the disruption effect on cytoskeletal structure of blebbistatin or nocodazole was relatively gentle, with partial cytoskeleton disruption, resulting in reduced amount and thickness of stress fibres and a 46% and 5% decrease in Young’s modulus, respectively.

In contrast, an increase in the average Young’s modulus of cells was observed with 60 min of colchicine treatment. However, no obvious change of the cytoskeletal structure was observed in the confocal images. The Young’s modulus data reveals regions of stiffer values within the cells while maintaining the initial cell morphology, with a 1.7-fold increase of the average cell stiffness.

These results agree with previous research that demonstrated the effect of these chemical treatments on intracellular structures,^17,40,41,43^ and additionally provided further information regarding the biomechanical properties of the hMSCs via the Q-AFM results. More subtle changes, such as those depicted in the cells treated with nocodazole or colchicine, were evident in the Young’s modulus maps thus demonstrating the sensitivity in this approach.

### Tracking cytoskeletal remodelling and biomechanical changes in real time

The ability of hMSCs to regenerate their cytoskeletal structure post-treatment was also of interest, hence we explored the temporal responsiveness of the cells using Q-AFM. Dynamic biomechanical changes were quantified by performing 20 min scans over a period of 3-4 h, in which the hMSCs were exposed to the chemical treatments discussed above and then subsequently replacing the media to allow the cells to recover (Fig 3), with the experimental timeline show in Fig 3a.

**Figure 3.**
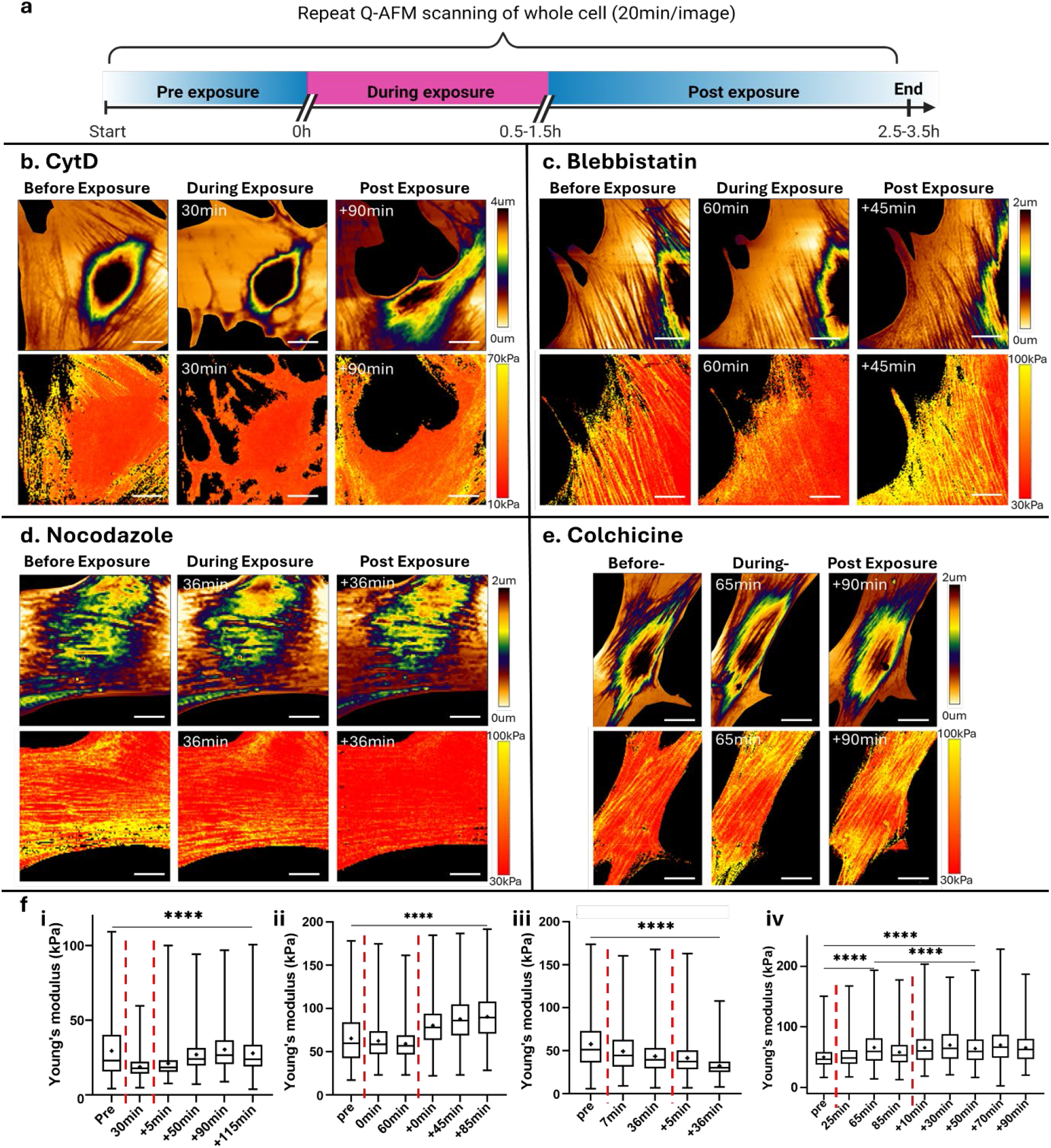
Representative AFM results of monitoring hMSCs with cytoskeleton disruptor treatment in real-time. (a) Experimental timeline, (b) 1 µg/mL CytD, (c) 12.5 µM blebbistain, (d) 5 µg/mL nocodazole and (e) 5 µM colchicine. Top rows represent AFM topographical images, and bottom rows represent the associated Young’s modulus maps. Scale bar is 20 µm. (f) The associated Young’s modulus distribution min-max box and whisker plot under each chemical treatment with i-iv corresponding to b-e. (Star shape and bar within box represents mean and median value respectively, bars outside box represent upper extreme and lower extreme, red dot lines highlight the timepoint of application and removal of disruptors). ‘Xmin’ represents x min during treatment, ‘+xmin’ represents x min after replacing fresh L15 media. **** indicates statistically significance as measured by Welch’s t test (P ≤ 0.0001).

The temporal response of the hMSCs treated with CytD (Fig 3a, 3i) showed a significant depolymerisation of the cytoskeleton after 30 min of treatment, followed by the rebuilding of the cytoskeleton network to return the average Young’s modulus of the cell to the pre-treatment values by 90 min post-exposure (initially 29 ± 17 kPa reduced 18 ± 6 kPa (P < 0.0001), and recovered to 30 ± 13 kPa), as shown in Fig 3c. The topographical scans show the expected morphological disruption of the cell as the actin was depolymerised, with 20% cell area shrinkage, although interestingly the presumed nuclei region remains intact as the cell remodels around it. The deformation of the cell morphology still apparent at 115 min post treatment is attributed to the AFM tip disturbing the cell as it remodelled.

The blebbistain treatment induced the initially thick cytoskeleton to soften as myosin II was inhibited, inferring that the cell lost contractility (Fig 3b). This effect was quantified by the Young’s modulus changes, showing significant cell softening from initially 54 ± 26 kPa down to 50 ± 16 kPa (Fig 3c), which could potentially lead to adipogenic differentiation^18,44^. With binding of myosin to actin inhibited by blebbistatin,^45^ further inhibiting actin activation of NMM II ATPase activity,^46^ the generation of cytoskeleton tension is restrained, which highly impacts processes such as cell motility, cytokinesis and differentiation direction.^47^ After the media change, the cell quickly remodelled the cytoskeleton, becoming demonstrably stiffer than the initial untreated state (74 ± 28 kPa) 45 min post-exposure. It was reported that subsequent washout of the blebbistatin could also allow full recovery of MyoD expression,^46^ meaning that the cells still possess their differentiation ability. Interestingly, comparing the treatment of CytD and blebbistatin, the effect of blebbistain on disrupting cytoskeletal structure was relatively gentle, as the binding target was different, with CytD fully depolymerising actin, while the effect of blebbistain on cell softening was stronger as cells tended to lose contractile force.

The nocodazole treatment (targeting the microtubule) had no observable impact on the cell morphology over the experiment, and there was no obvious change in the cytoskeletal structure in the topographical scans, inferring that the microtubule does not primarily maintain cytoskeleton network and support cell shape, unlike the actin filament. However, the Young’s modulus of the cell decreased significantly during the treatment (57 ± 28 kPa to 49 ± 23 kPa) and continued to decrease once the media was replaced (32 ± 10 kPa at 90 min post-exposure) (Fig 3iii). In previous studies, treating cells with 1µg/mL nocodazole in osteogenic media resulted in complete disruption of microtubule arrangement, while calcium deposition exhibited same level as untreated cells.^48^ On the contrary, with much lower concentrated nocodazole 50 nM (∼ 0.15 µg/mL) in basal media, hMSCs presented halved mineralisation compared to the control group (treated with DMSO, solvent of nocodazole),^49^ indicating microtubule cytoskeleton contributes toward osteogenic differentiation, while with osteogenic supplements, the does-dependent disruption of microtubule by nocodazole has limited effect on cell fate control.

The colchicine treatment, also targeting microtubules resulted in an increase in cell stiffness (50 ± 17 kPa to 55 ± 21 kPa), with the increase in actin structures clearly observable over the treatment period and beyond (Fig 3e), agreeing with colchicine effect of inhibiting microtubule polymerisation while promoting actin filament formation. With 85 min treatment, the average Young’s modulus of the cell (67 ± 27 kPa) was significantly stiffer than the initial measurement, remaining at this average stiffness 90 min post-treatment, Fig 3iv. The conversion of hMSCs into neuron-like cells was restricted with 1 µg/ml colchicine.^50^ While the stiffening of hMSCs has higher chance to differentiate into osteocytes. The effect of colchicine on osteogenic gene expression has not been addressed yet.

With the same cells imaged under chemical treatments at different time point shown in Fig 3, at 20 min per frame, the morphological changes and biomechanical changes in the cells were observed, providing cell structure and mechanics images with much higher resolution in a short time frame comparing to the traditional force-volume mapping.^51^ For example, the cell area shrank 20% with CytD treatment due to the cytoskeletal structure disruption and remained similar size even 90 min post exposure while inner cytoskeleton network was rebuilt during cell recovery. The consecutive images of all cells over the experiment are shown in Fig S3, with 6, 7, 5 and 14 total scans, including pre-, during-, post-exposure on the same cells for CytD, blebbistatin, nocodazole and colchicine treatment respectively, depending on the speed and extent of cell response to the applied chemicals.

### Intracellular features characterisation during exposure

The structure of actin fibres, which are a major component of the cytoskeleton, is a primary contributor to the quantifiable stiffness of the cell.^47,52^ Mapping changes in these intracellular features is difficult, and monitoring dynamic remodelling of individual features is important to fully understand the contribution of these intracellular features to cell stiffness during exposure. Line profiles of a single stress fibre, across multiple stress fibres, and across the cell without visible fibres were extracted and shown in Fig 4.

**Figure 4.**
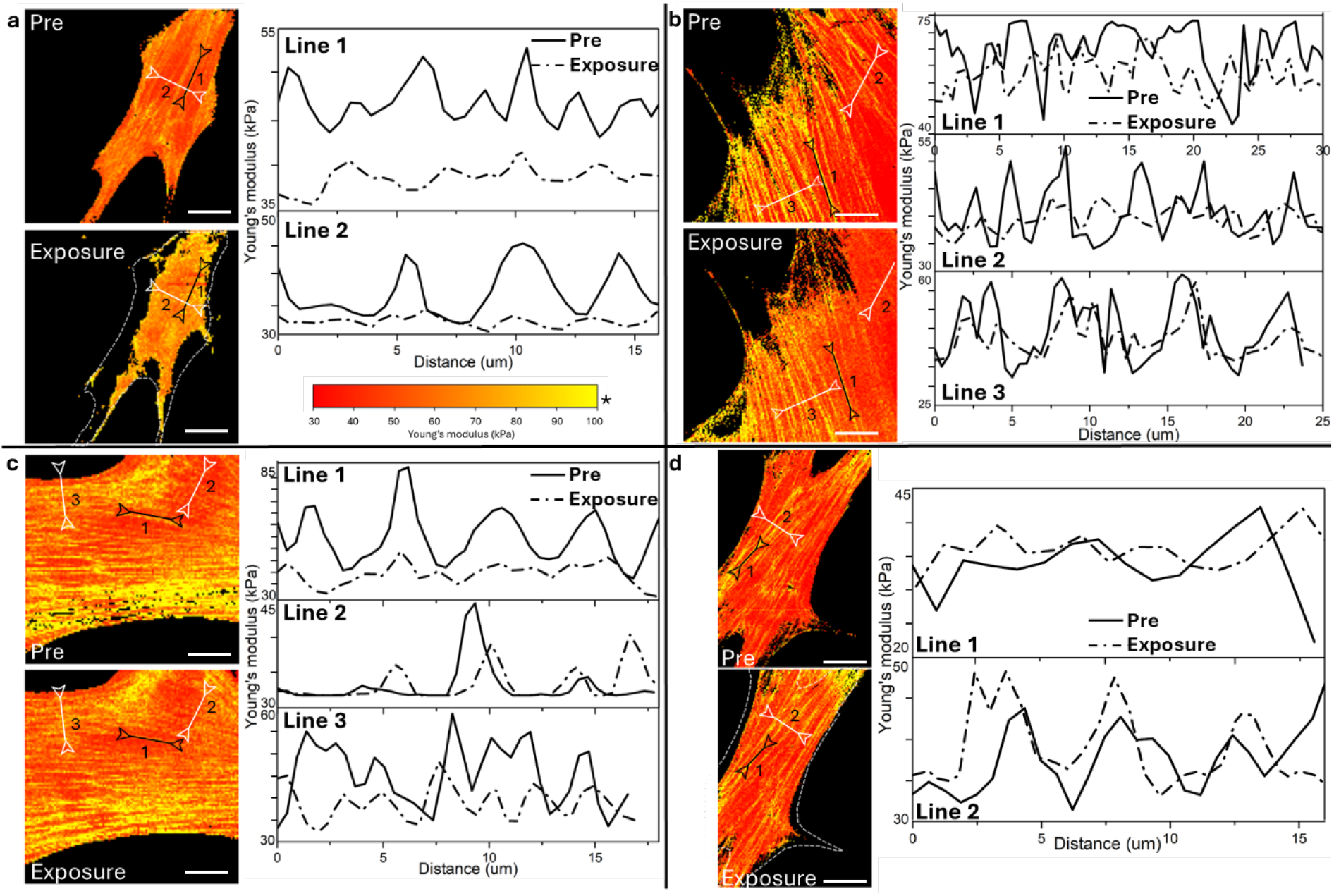
Representative analysis of intracellular features (single stress fibre) dynamics of hMSCs under (a) 1 µg/mL CytD, (b) 12.5 µM blebbistain, (c) 5 µg/mL nocodazole, and (d) 5 µM colchicine treatment. The solid lines and dash dot lines represent the corresponding profile of pre-treatment and during-exposure cells respectively indicated in the images. The dot lines shown in **a**- and **d**-Exposure outline cell shape before chemical treatment. Scale bar is 20 µm. *Colour bar 30-100 kPa applied to all images except **a**-Exposure with range 10-30 kPa and **c**-Exposure with range 10-80 kPa.

Depolymerisation of actin in the cytoskeleton leads to cell softening; this is clearly apparent with the CytD treatment, shown in Fig 4a, where all visible fibre structures were disrupted and the cell morphology changed dramatically, shrinking after treatment. Changes to individual intracellular features was quantifiable, where the Young’s modulus along a clear actin fibre structure changes significantly post-treatment (Fig 4a, line 1). The lateral features along the fibre are still visible, however the average Young’s modulus value along the fibre has clearly reduced (from 48 to 39 kPa) concomitantly. A cross-section across several actin fibres (Fig 4a, line 2) demonstrates the significant change in Young’s modulus with high spatial resolution as the fibres undergo depolymerisation.

The individual actin structures dissipated with the addition of blebbistatin, shown in Fig 4b, contributing to the overall reduction in the average Young’s modulus of the cell. The effect of the myosin II inhibition to soften actin structures is clear when analysing the individual structures; along a single feature (Fig 3b, Line 1) the Young’s modulus has reduced on average 10%.^53,54^ The thickness of actin bundles was 1.3-fold thicker than that after blebbistain application, which might also affect cell stiffness. Spatially quantifying the Young’s modulus crossing multiple stress fibres, (Fig 3b, Line 2 and line 3), clear peaks of Young’s modulus were observed, corresponding to the specific stress fibre, presenting general softening of single stress fibre. Comparing the profiles of line 2 and line 3, the actin features react heterogeneously, with originally weaker actin softening to a higher degree.

Similar results of measuring single stress fibres under nocodazole treatment was observed,^55^ Fig 3c, line 1, showing an overall Young’s modulus decrease as nocodazole inhibits microtubule polymerisation. Crossing multiple actin bundles (Fig 4c line 3), all stress fibres became softer after 36 min nocodazole treatment, while the fibre hadn’t been totally depolymerised. Measuring the region around line 2 without clear stress fibres, the baseline of Young’s modulus before and during the treatment collated, showing that under nocodazole treatment, the decreasing stiffness and amount of stress fibre was the main factor of cell softening, while the other cell part remained mostly unchanged of stiffness.

With 25 min colchicine exposure, the single actin fibre stiffness remained unchanged, while the actin bundle became thicker as actin filament polymerisation was promoted, resulting in overall cell stiffening.

## Discussion

The interplay between cytoskeletal structure, biomechanical properties, and stem cell fate are key to using external cues to direct differentiation. It has been well reported that hMSC differentiation can be controlled through manipulation of the cytoskeleton, through either chemical and physical cues,^17,56,57^ which in turn modulates intracellular tension. With mechanical stimulus, the mechanotransduction pathways are activated, leading to fate decision by the stem cells. Specifically, sensing the external stimulus via plasma membrane, involving integrins, lipid rafts, ion channels and F-actin, the physical signal is converted into a biochemical signal to induce intracellular change and transmitted through cytoskeleton network, resulting in cytoskeleton reorganization, cell shape restructuring and cell mechanics dynamics. Those eventually affects nuclear properties, YAP/TAZ translocation and gene expression, ultimately regulating stem cell fate.^19,20^

Hence, the importance of how the cytoskeleton remodels and changes in response to its surrounding environment is clear when understanding cellular response. However, performing real-time studies on living cells has historically been highly challenging. Biosensors such as FRET or SiR-actin can reveal highly resolved and sensitive information about intracellular tension or actin dynamics.^58,59^ Linking the cytoskeletal remodelling to quantifiable biomechanical data with no transfection, biosensors, or staining however offers many advantages. Foremost that Q-AFM can be applied to an adherent cell line on practically any type of material or substrate.

Regulating the cytoskeleton structure and cell stiffness could affect cell fate decision.^46^ hMSCs with depolymerised cytoskeleton induced by CytD treatment, were reported to favour adipogenesis and show decrease osteogenesis.^60^ With Q-AFM monitoring, significant cell softening was observed during CytD exposure, inferring decreasing in cell stiffness elicits adipogenesis, which is supported by the results that cell softening and stiffening was generally observed with growth factor-induced adipogenesis and osteogenesis respectively.^18,44^ Removing the CytD, hMSCs recovered their cytoskeleton network with increasing cell stiffness. However, cells may still undergo adipogenesis with cell’s memory, presenting a much lower-level of alkaline phosphatase activity, despite the recovery in cell shape and actin cytoskeleton organisation (48 h CytD treatment and 24 h recovery).^48^

Here we at first demonstrated the cellular structure, and how cell morphology measured by Q-AFM is consistent with confocal images, while the identical features presented in cell stiffness maps also correlated to the topographical images. With force-volume mapping mode, cell morphology and biological structures had been accurately co-located via confocal microscopy with the same structures in AFM by comparing their 3D projections.^61^ The measured features of cells via Q-AFM mode were mainly cytoskeleton network, which was confirmed by Z-stack confocal images of the same cells stained with actin dyes, shown in Fig 1. Apart from showing intracellular structures, Young’s modulus maps also revealed cell stiffness distribution within the cells, linking features reorganisation to stiffness dynamics.

Secondly, we have directly characterised the cytoskeletal remodelling of hMSC under four different cytoskeletal disruptors, using Q-AFM to observe real-time changes and quantified the Young’s modulus to a high spatial resolution. Compared to current literature, we have presented the first study using Q-AFM to map live cells at a far greater resolution than typically shown, in addition to the temporal analysis.

During exposure to the chemicals, cytoskeleton depolymerisation/polymerisation to different extents was monitored and the cell mechanics were quantified. Specially, with CytD treatment, the cytoskeleton was significantly disrupted, which agrees with the fluorescent images reported in Fig 2 and also fluorescent results in previous papers.^17^ The recovery of cytoskeleton network after replacing fresh media, also observed by Gregory Yourek et al., leads to increasing cell Young’s modulus, which hadn’t been reported before. Cell response on the other three chemical treatments presented here also corresponded to the results presented in literatures,^40,41,43^ while more details of cell mechanics from the same cells in real-time were achieved.

Cell mechanics changes during differentiation, apoptosis and other stimulation treatment have been measured with traditional AFM. However, only end point measurements were mainly conducted, without monitoring the same single cells change in real-time, which leads to unknown mechanisms of hMSCs responding to external factors. Moreover, it is hypothesised that stem cell mechanics, specifically Young’s modulus could be an early marker to predict cell fate,^62,63^ which reveals the importance of characterisation cell stiffness under external stimuli continuously.

The Q-AFM is powerful for characterising physical and mechanical properties of any adherent cell types, enabling the examination of how new materials impact cells. For instance, by directly seeding cells on materials with diverse features like topography (e.g., star shape, flower shape)^64^, patterning (e.g., micropillars arranged in grid platform, microgrooves)^65,66^ and stiffness (ranging from 0.1 to 40 kPa)^16^, stress fibre distribution and reorganisation as well as intricate cell mechanics of the same cells during adaption to the substrate could be explored with Q-AFM measurements. Additionally, Q-AFM allows real-time monitoring of cell behaviour when culturing cells on/with materials such as mesoporous silica nanoparticle carrying chemicals that release drugs (e.g., cytoskeleton disruptors, growth factors) at controlled rates, influencing cell fate.^67^ Notably, Q-AFM offers greater flexibility for studying cell interactions with non-transparent materials since AFM measurements rely on physical contact with cells rather than light or laser paths. This approach aids in the understanding of cell-material cinteractions and the impacts of material properties on cell behaviour and fate regulation, without requiring modifications to either the cells or the material surfaces.

Furthermore, the Q-AFM measurement system could be easily coupled with an external platform/device to monitor cellular dynamics with external stimulation, helping reveal the mechanism of stem cell response to dynamic factors, eventually achieving precise cell fate regulation for tissue engineering application. By integrating Q-AFM with a uniaxial motorised cell stretching device,^68^ cellular response of stretching deformation on living cells can be identified to study mechanotransduction processes. Adding on a surface acoustic wave stimulation platform to apply shear stress to stem cells,^69^ cytoskeleton realignment, cell stiffness dynamics during and post treatment can be used to study the mechanism of triggering osteogenesis, in turn to guide the applied parameters. Apart from cell stiffness, cell adhesion can also be characterised with Q-AFM to understand cell-material interactions, and cell-cell communication under external treatment.

## Conclusion

In this paper, we used Q-AFM to measure the cellular structure and cell mechanics of hMSCs under four different cytoskeleton disruptor treatments in real-time. The increasing or decreasing cell Young’s moduli were monitored along with intracellular features and cell morphology dynamics observation, marking Q-AFM as a powerful technique to achieve temporal analysis of cell mechanics with external stimulus. Measuring cell morphology, stiffness, and adhesion by Q-AFM with high resolution at fast speeds compared to traditional force-volume mapping mode, allows real-time measurements of cell mechanics in detail, correlating dynamic cell structure and cell mechanical properties. With this technique, cell mechanotransduction mechanisms could be further studied, which would highly contribute to precise hMSCs fate regulation for tissue engineering applications under external physical stimulation.

## Author Contribution

KZ: Investigation, methodology, formal analysis, visualisation, writing – original draft preparation (lead). CR: Investigation, writing – review & editing (supporting). JB: Software. KF: Funding acquisition, supervision. AE: Conceptualization (equal), supervision, resources, writing – review & editing. AG: Conceptualization (equal), supervision, funding acquisition, resources, methodology, writing – review & editing (lead).

## Acknowledgement

This research was funded by the Australian Government through the Australian Research Council by Discovery Project (DP200100612).

## Notes

### Competing Interest Statement

The authors have declared no competing interest.

