## Supplementary Data for "Real-time Biomechanical Characterisation of Cytoskeletal Remodelling"

### S1: Comparing substrate uncorrected and corrected Young's modulus fitting

Our customised Python programming includes various versions of equations based on the basic Hertz model (Eq. 1), tailored to meet specific project requirements. Notably, for a parabolic sharp tip, the corrected equation developed by Garcia, P. D., & Garcia, R. (Eq. 2) is commonly employed to minimise substrate effects. In our analysis, we compared the fitting results of Young's modulus obtained from the substrate uncorrected and corrected models.

$$F = \frac{4}{3}\frac{E}{1-\upsilon^2}\sqrt{R}\delta^{3/2} \qquad (1)\ [29]$$

$$F = \frac{16}{9\pi}E\sqrt{R}\delta^{3/2}\left[\frac{1}{h^0} + \frac{1.133\sqrt{\delta R}}{h} + \frac{1.497\delta R}{h^2} + \frac{1.469\delta R\sqrt{\delta R}}{h^3} + \frac{0.755\delta^2 R^2}{h^4}\right] \qquad (2)\ [32]$$

Where F is force, $\delta$ is cell indentation , R is probe tip radius, $\upsilon$ is Poisson's ratio, h is height of the sample at that location, and E is Young's modulus.

Referring to Fig S1b and S1c, we observed no significant difference between the results obtained from these two models. Both exhibited high resolution and consistent stress fibre structures. While the actual Young's modulus value of the uncorrected model is 7% higher than that obtained from the corrected model, the distribution of Young's modulus was similar (Fig S1d). This infers that with our fitting parameter (fitting depth less than 18% of cell height), substrate effect had already been highly limited. Consequently, both versions of the Young's modulus fitting are deemed suitable for data processing. It is imperative to maintain consistency by using the same version of the model when processing data from the same cell at different time points throughout the experiment. This ensures coherence and comparability of results across experiments.

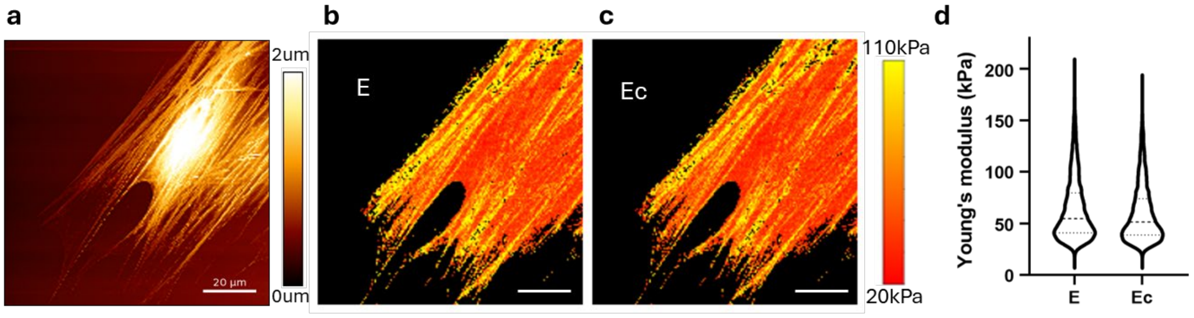


Figure S1. Comparing Young's modulus obtained with different fitting equations. (a) Topographical image, (b) Young's modulus map with Hertz model fitting, (c) Young's modulus map with substrate corrected Hertz model fitting. (d) Young's modulus distribution. Scale bar is 20 µm.

S2 preliminary experiments to determine the suitable concentrations of each chemical.

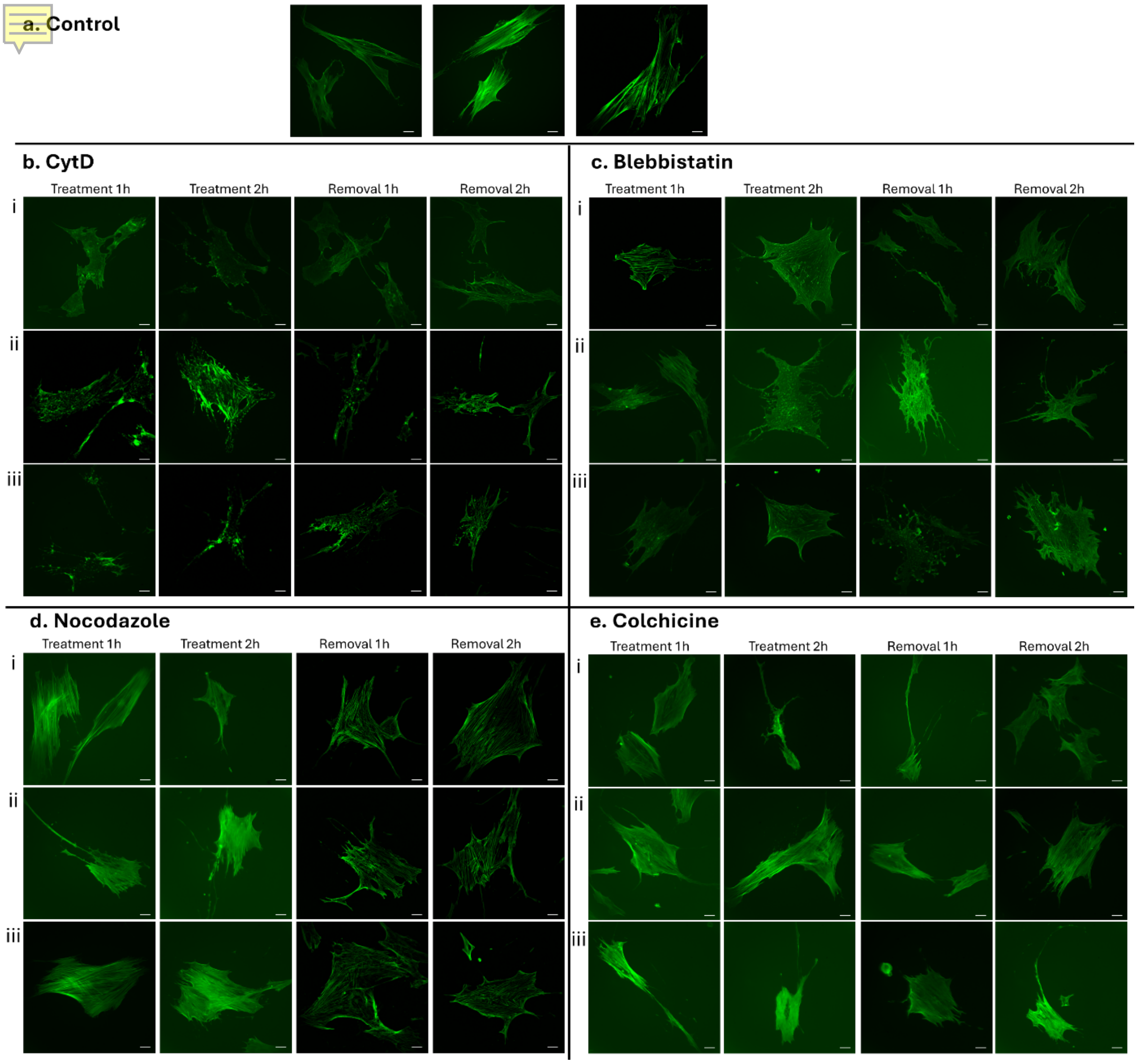


Figure S2. Widefield images of hMSCs with treatment of cytoskeleton disruption chemicals in different concentrations. (a) Control group, (b) CytD with 0.1 μg/mL (row i), 0.5 μg/mL (row ii), 1.0 μg/mL (row iii), (c) Blebbistatin with 12.5 μM (row i), 25 μM (row ii), 50 μM (row iii), (d) Nocodazole 1 μg/mL (row i), 5 μg/mL (row ii), 10 μg/mL(row iii), (e) Colchicine 0.5 μM (row i), 5 μM (row ii), 50 μM (row iii). Scale bar is 20 μm.

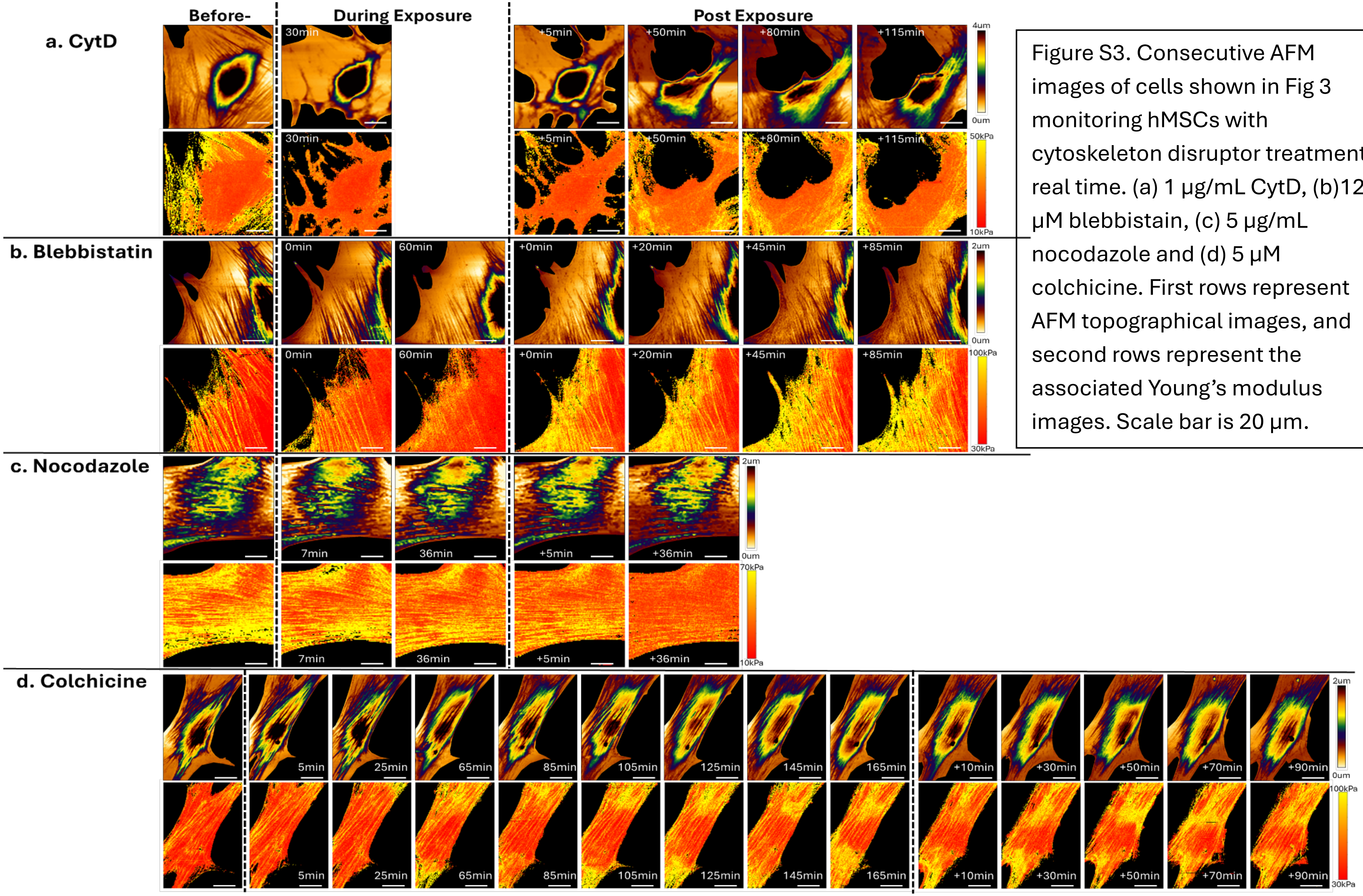


Figure S3. Consecutive AFM images of cells shown in Fig 3 monitoring hMSCs with cytoskeleton disruptor treatment in real time. (a) 1 μg/mL CytD, (b)12.5 μM blebbistain, (c) 5 μg/mL nocodazole and (d) 5 μM colchicine. First rows represent AFM topographical images, and second rows represent the associated Young's modulus images. Scale bar is 20 μm.
